# Unraveling the world of halophilic and halotolerant bacteria in cheese by combining cultural, genomic and metagenomic approaches

**DOI:** 10.1101/2020.11.03.353524

**Authors:** Caroline Isabel Kothe, Alexander Bolotin, Bochra-Farah Kraïem, Bedis Dridi, FoodMicrobiome Team, Pierre Renault

## Abstract

Halophilic and halotolerant bacteria are generally assumed to live in natural environments, although they may also be found in foods such as cheese and seafood. These salt-loving bacteria have only been occasionally characterized in cheese, and studies on their ecological and technological functions are still scarce. We therefore selected 13 traditional cheeses in order to systematically characterize these microorganisms in their rinds via cultural, genomic and metagenomic methods. Using different salt-based media, we identified 35 strains with unique 16S rRNA and *rpoB* gene sequences, whose whole genome was sequenced. The most frequently isolated species are the halotolerant Gram-positive bacteria *Brevibacterium aurantiacum* (6) and *Staphylococcus equorum* (3), which are also frequently added as starters. Their genomic analyses confirm the high genetic diversity of *B. aurantiacum* and reveal the presence of two subspecies in *S. equorum*, as well as the genetic proximity of several cheese strains to bovine isolates. Additionally, we isolated 15 Gram-negative strains, potentially defining ten new species of halophilic cheese bacteria, in particular for the genera *Halomonas* and *Psychrobacter*. The use of these genomes as a reference to complement those existing in the databases allowed us to study the representativeness of 66 species of halophilic and halotolerant bacteria in 74 cheese rind metagenomes. The Gram-negative species are particularly abundant in a wide variety of cheeses with high moisture, such as washed-rind cheeses. Finally, analyses of co-occurrences reveal assemblies, including the frequent coexistence of several species of the same genus, forming moderately complex ecosystems with functional redundancies that probably ensure stable cheese development.

**IMPORTANCE:** Salt is commonly added to food to avoid the growth of pathogens by lowering water activity, resulting in profound changes in the medium that lead to the development of particular ecosystems dominated by halophilic and halotolerant bacteria, communities that probably originate in the natural environment. In order to explore these communities that have been poorly studied in food up until now, we developed a combined approach that includes cultures, genomics and metagenomics to deconstruct these ecosystems in cheese rinds. This approach allowed us to isolate 26 different species, ten of which belong to still undescribed species that could be used as references to promote advances in functional studies of this particular world. The metagenomic scan of 74 cheese rind samples for the assembly of 66 halophilic and halotolerant species showed that these bacteria are widely distributed and form moderately complex ecosystems where related species coexist and probably jointly contribute to safe and efficient cheese development.

## Introduction

Occurring naturally in nature, salt has been harvested since ancient times on the shores of lakes, seacoasts and oases. There is even evidence that it was produced industrially during protohistory, in the 5-6^th^ millenium BC [1–3]. Regardless of its source (rock salt, sea salt, spiced salt, etc.), raw salt serves as a flavor enhancer and increases the palatability of food, while allowing the preservation of certain foodstuffs, especially meat and fish.

Salt is effective as a preservative because it reduces water activity in foods [4, 5]. Its addition can reduce the rate of microbial growth in foods due to osmotic shock and other disorders that interfere with cell enzymes or that expend energy to exclude sodium ions from the microbial cell [6]. Although high levels of salt totally inhibit the growth of most microorganisms, moderate levels create an inhospitable environment for the majority of pathogens and promote the growth of certain microorganisms in various food products. Despite advances in food processing, storage, packaging and transportation have largely diminished this role, and salt is still widely used as one of the multiple means to control food safety. Moreover, it frequently plays a central role in the production of fermented foods (i.e., pickles, sauerkraut, cheeses, Asian seafood, fermented meats, etc.), not only contributing to an extended shelf life, but also leading to the development of particular aroma, texture, nutritional and beneficial health properties, eventually becoming part of the cultural patrimony in many countries. Whereas many studies have focused on lactic fermentation in recent decades, the interest of the salt-driven development of non-pathogens in fermented food is more recent and at its beginning stages.

Microorganisms that grow in the presence of salt may be subdivided into two categories: (1) halotolerants, which are able to grow in the presence or absence of salt; and (2) halophiles, which require salt to develop [7, 8]. The precise definition of a halophile diverges depending on the authors, some of whom include any organism that requires percentages of around 3.5% of salt, as in seawater [7, 9], while others consider only those that grow optimally at 5% or above, and tolerate at least 10% of salt [10]. Such halophilic bacteria have been widely described and isolated in different natural ecosystems such as soil, salt lakes and seas, and are of interest as producers of pigments and antibacterial activities [11, 12], as well as a variety of bioactive compounds [13, 14]. Halophilic and halotolerant bacteria have also been isolated from foods such as salt meat, shrimp, fermented fish sauce, sea food, poultry and cheese [15–18]. Several strains found in these foods have already been characterized, but their functions are still unknown. Certain studies consider that halophilic bacteria are involved in food spoilage and are thus considered to be undesirable in food processing environments [16, 17, 19]. Other authors report that several species produce significant quantities of volatile compounds such as sulfides, acetone, ammonia and ethanol, suggesting that they have a potential function in aroma production [15, 20].

Overall, studies on halophilic and halotolerant bacteria in food are still scarce. A comprehensive overview of all ecosystem agents is therefore necessary in order to understand their functions, manage their evolution in the process and, eventually, understand the association of these microorganisms with the shelf life of food products. Cheese is part of these ancient fermented foods in which salt addition may lead to major changes in its processing. There are more than 1,000 distinct cheese types worldwide, with a variety of textures, appearances, aromas and flavors that can be attributed to the technological development of complex and specific microbial communities, as well as to local factors such as milk source and farming practices [21–23]. The salting process might thus be one of the factors that strongly influence the way a microbial community will develop in cheese. Salt can be added in different ways - in crystal form or in solution, by brining or applying brine directly to the curd, by rubbing, smearing or scraping the surface, with clear salt solutions or historical old brines - and each of these processes may be applied once or several times and at different time scales (**Fig. 1**). Indeed, recent studies based on non-cultivable methods have shown that halophilic bacteria may be the dominant microbiota on cheese rinds, suggesting that their role has been underestimated until now [24, 25]. A better understanding of their origin and how these organisms evolve during the different stages of cheese development could be of major interest to determine the role of salt in cheese ecosystem organization.

**Figure 1.**
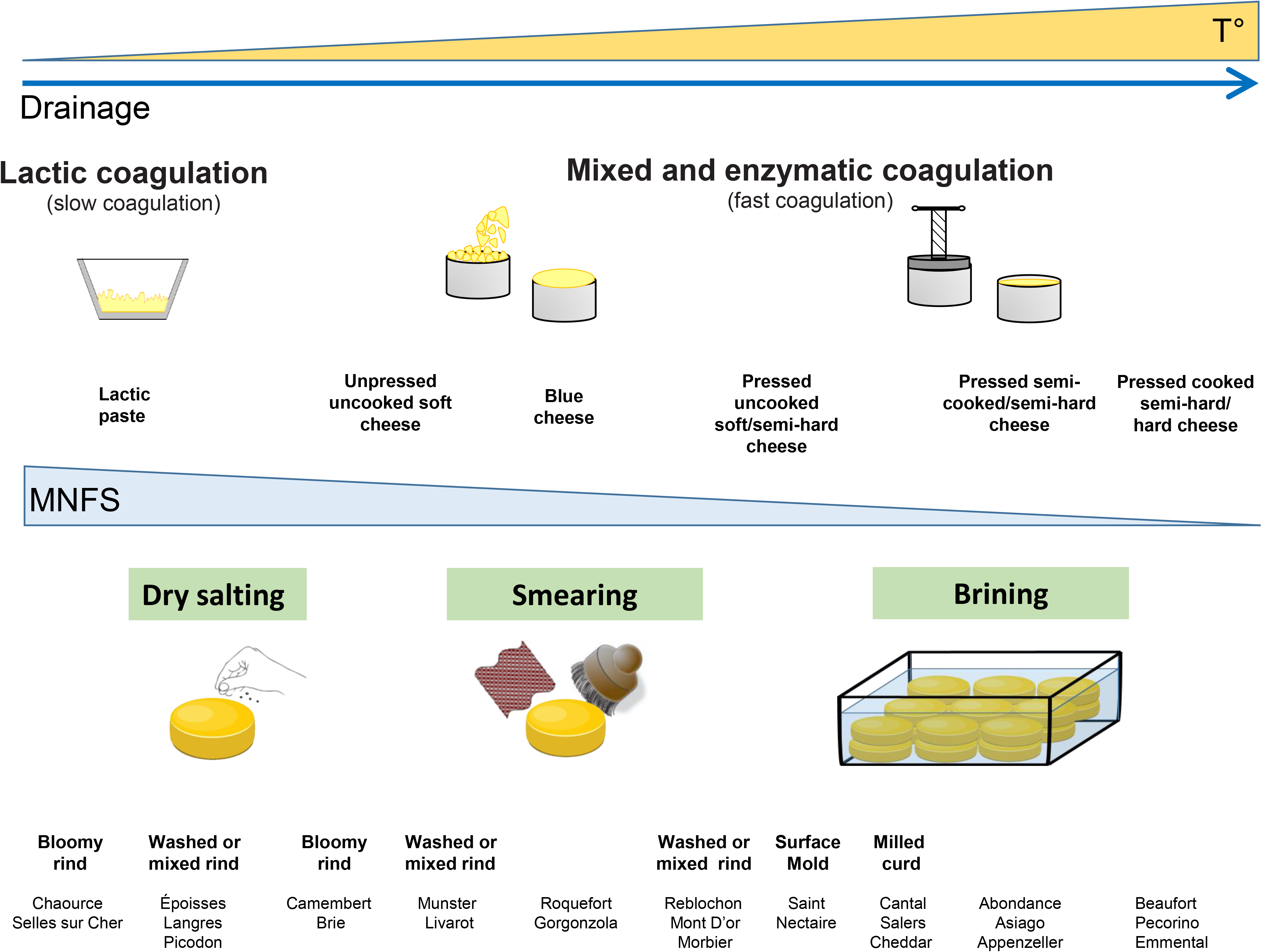
Cheese processing including examples of the salting process. Processes were classified as proposed by Almena-Aliste & Mietton [40]. The effect of water activity depending on the process, in particular, on drainage and temperature, is primarily dependent on the moisture in non-fat substances (MNFS), whereas the salting level decreases this effect. Examples of cheeses are given to illustrate this figure.

We therefore decided to enlarge the data repository of these salt-tolerant and salt-dependant bacteria in cheese in order to determine and produce a precise overview of their presence in cheeses produced using different technologies. For this purpose, we used culture methods to isolate and identify - in a systematic way - halophilic and halotolerant bacteria in 13 artisanal cheese rinds produced by different salting processes. We then combined genomic and metagenomic approaches that revealed potential new species, a wide diversity of halophilic and halotolerant bacteria in cheese rinds, and the coexistence of these species in this food ecosystem.

## Results

### Abundance of halophilic and halotolerant bacteria in cheese rinds

For this study, we selected 13 artisanal cheese rinds whose main technological features are presented in **Table 1**, in order to systematically characterize their halophilic and halotolerant microorganisms by culture methods. Different cheese technologies were included: lactic paste, unpressed or pressed, uncooked or cooked, soft or semi-hard and blue cheeses. Moreover, the cheeses involved different salting processes such as brining, dry salting, rind washing or smearing.

**Table 1:**
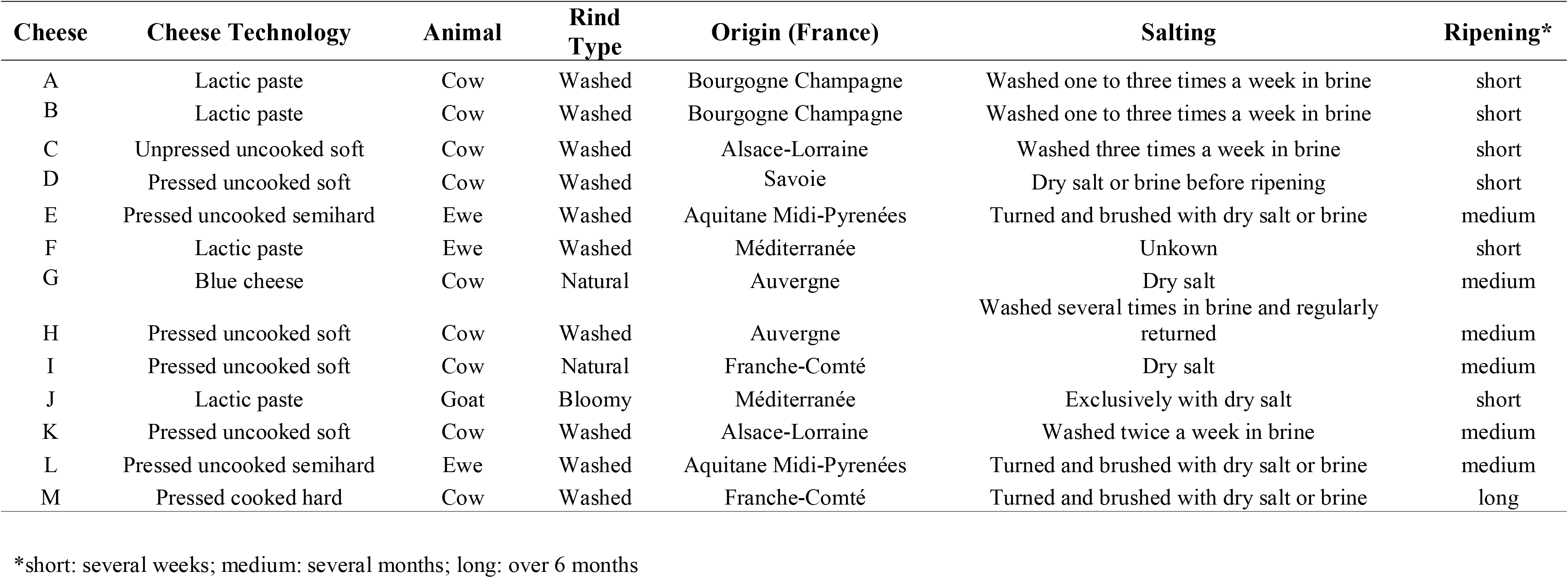
Metadata describing the 13 cheese rind samples.

| **Cheese** | **Cheese Technology** | **Animal** | **Rind Type** | **Origin (France)** | **Salting** | **Ripening\*** |
| --- | --- | --- | --- | --- | --- | --- |
| A | Lactic paste | Cow | Washed | Bourgogne Champagne | Washed one to three times a week in brine | short |
| B | Lactic paste | Cow | Washed | Bourgogne Champagne | Washed one to three times a week in brine | short |
| C | Unpressed uncooked soft | Cow | Washed | Alsace-Lorraine | Washed three times a week in brine | short |
| D | Pressed uncooked soft | Cow | Washed | Savoie | Dry salt or brine before ripening | short |
| E | Pressed uncooked semihard | Ewe | Washed | Aquitane Midi-Pyrenées | Turned and brushed with dry salt or brine | medium |
| F | Lactic paste | Ewe | Washed | Méditerranée | Unkown | short |
| G | Blue cheese | Cow | Natural | Auvergne | Dry salt | medium |
| H | Pressed uncooked soft | Cow | Washed | Auvergne | Washed several times in brine and regularly returned | medium |
| I | Pressed uncooked soft | Cow | Natural | Franche-Comté | Dry salt | medium |
| J | Lactic paste | Goat | Bloomy | Méditerranée | Exclusively with dry salt | short |
| K | Pressed uncooked soft | Cow | Washed | Alsace-Lorraine | Washed twice a week in brine | medium |
| L | Pressed uncooked semihard | Ewe | Washed | Aquitane Midi-Pyrenées | Turned and brushed with dry salt or brine | medium |
| M | Pressed cooked hard | Cow | Washed | Franche-Comté | Turned and brushed with dry salt or brine | long |

\*short: several weeks; medium: several months; long: over 6 months

The halophilic and halotolerant bacterial population level was estimated by plate counts on several media and temperatures in order to optimize their growth and maximize the potential to isolate diverse strains. Overall, our results demonstrated the presence of dense bacterial populations (around 7-8 log CFU/g) on the surface of most cheese rind samples, with a minimum of 4.0 and a maximum of 9.5 log CFU/g (**Figure S1**). No major differences in bacterial counts were found between LH and MB media, whereas lower counts were obtained with HM medium for most cheeses, in particular Cheeses A, B and D. The effect of increasing salt was tested on MB medium and its increase negatively affected bacterial counts, in particular for Cheeses A, B and F. Interestingly, counts in MB+8% salt do not present major differences with those obtained in HM. The lower counts obtained in these two media, compared with other ones, could be due to a higher salt content: 8-10%, vs. 1 and 2% in LH and MB, respectively. Finally, the counts were similar at the three incubation temperatures tested (20, 25 and 30°C).

### Defining potential new food halophilic species

Two marker genes, 16S rRNA and *rpoB*, were used to identify and differentiate the bacterial isolates. For each cheese, representative isolates differing in at least one of these markers were selected for further study, and those grown on MB or its derivative containing higher salt content were preferentially chosen in the case of the identity of both markers. From ~320 isolates, we finally obtained 35 strains tentatively assigned to species, as described in **Additional File 1** and presented in **Table S1**. Of these strains, 20 belong to the Gram-positive group and 15 to the Gram-negative group, 17 and three isolates of which are assigned at the species level, respectively. To obtain reliable taxonomic data, we determined the genomic sequences of the 35 strains (**Table S2**) and performed an ANI analysis with closely related strains, as shown in **Table S3**.

Genomic analyses confirmed the assignation of the 17 Gram-positive bacteria carried out by the marker genes, while three remaining isolates belong to undescribed species (**Table 2**). *Brachybacterium* strain FME24 displays an ANI value of 83.17% with its closest relative *B. tyrofermentans*, indicating that it belongs to a new species. Similar analysis of *Brevibacterium* strains FME17 and FME37 showed that these two strains and *Brevibacterium sp.* 239c share an ANI of over 97% of each other (**Table S3**), but less than 87% with the closest reference species, indicating that they belong to a new species.

**Table 2.**
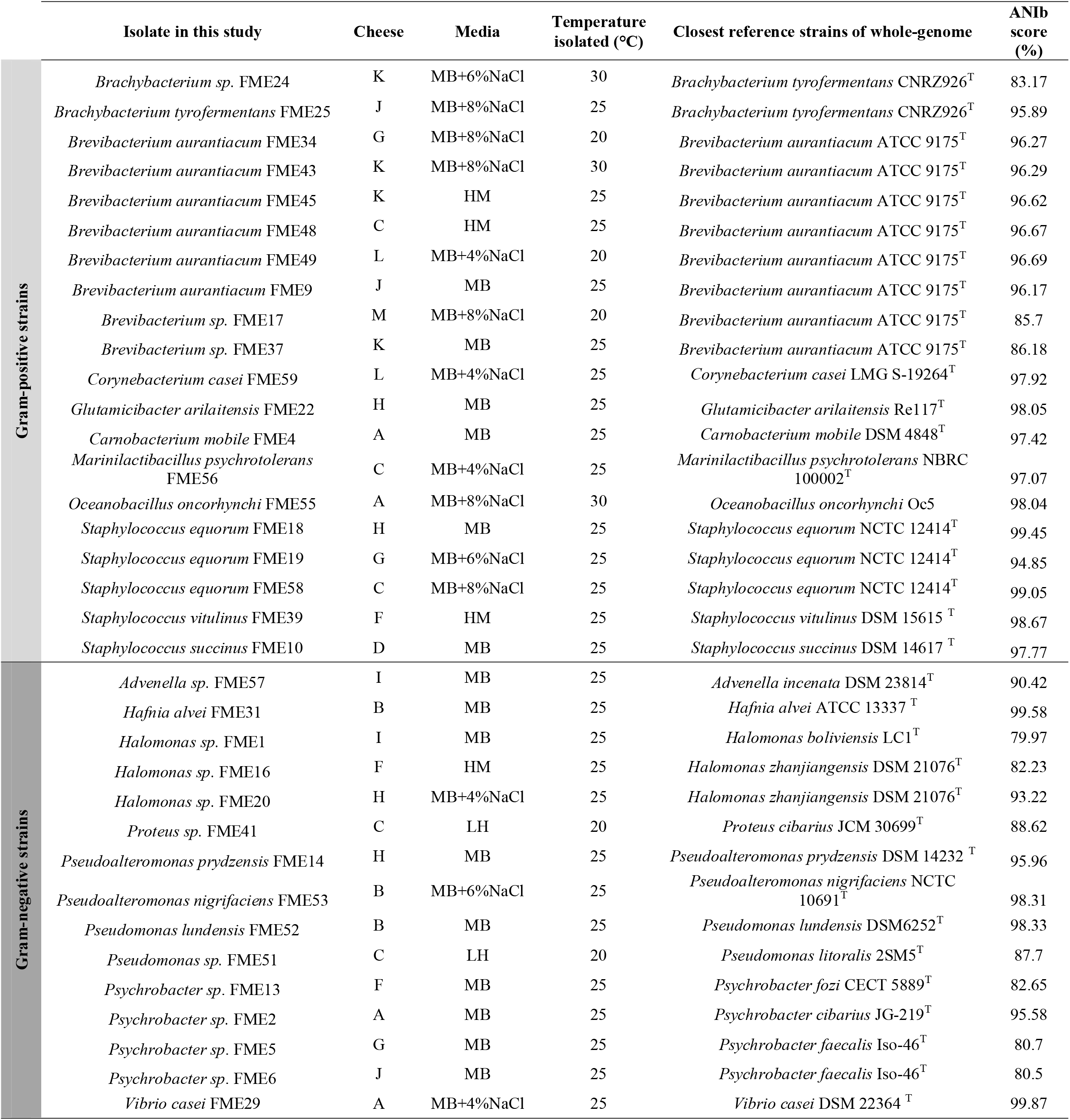
The 35 isolates and their cheese origin, media and temperature of isolation, as well as the closest species with their respective ANI values.

Concerning Gram-negative species, ANI analysis could assign only five isolates, while ten remained ambiguously or not assigned (**Table 2**). *Advenella sp.* FME57, isolated in this study, and *Advenella sp.* 3T5F (formerly referred to as *kashmirensis*) appear to belong to the same species (ANI=98.57%) but significantly differ from the *A. kashmirensis* type strain (WT001^T^) with which they share an ANI < 90% (**Table S3**). These two strains should therefore belong to a new species. Concerning the *Halomonas* genus, ANI analysis showed that none of the strains isolated here could be assigned to an already described species, including the FME20 strain whose 16S rRNA and *rpoB* analyses suggested its assignation to *H. zhanjiangensis*. Indeed, their ANI of 93.22% is below the threshold of 95% (**Table 2**). Results of marker analyses of the two *Pseudoalteromonas* strains (FME14 and FME53) remained ambiguous due to multiple hits with similar identities with different species (**Additional File 1**). The ANI analysis demonstrated that the FME14 strain could be assigned to *P. prydzensis* (ANI=95.96%) and FME53 to *P. nigrifaciens* (best score ANI=98.31%) (**Table 2**). Futhermore, *Proteus sp.* FME41 belongs to a new species since it shares only 88.62% ANI with its close relative *P. cibarius* JCM 30699^T^. Similarly, *Pseudomonas* strain FME51 does not belong to *Pseudomonas litoralis* since it shares an ANI of only 87.70% (**Table 2**). Regarding the *Psychrobacter* genus, FME2 shares ANI~95% with five strains of *Psychrobacter,* including *P. immobilis* and *P. cibarius* type strains (**Table S3**), leaving its assignation unresolved. Finally, *Psychrobacter* strains FME5, FME6 and FME13 could not be assigned to any already known species by both markers and ANI analyses (**Table S1**and **Table S3**). Since FME5 and FME6 strains display an ANI of 98%, these three strains may represent two new species.

Therefore, from these 35 isolates, we obtained strains belonging to 26 different species, ten of which potentially belong to new species: two to the Gram-positive group and eight to the Gram-negative group.

### Genomic diversity of *B. aurantiacum* and *S. equorum*

Among the Gram-positive bacteria, we isolated six *B. aurantiacum* strains from five different cheeses (**Table 2**), which, in addition to the 19 already sequenced genomes available in the NCBI database (**Table S4**), gave a total of 25 genomes. ANI analysis shows that they share over 97% identity and clustering analysis indicates the presence of four groups (indicated as A, B, C and D; **Fig. 2A**). To further determine their genetic diversity, we performed a pan-genome analysis that revealed an open pan-genome for 25 strains of *B. aurantiacum,* with a total of 10,823 genes (**Fig. S2A)**. We also analyzed the number of genes present in different numbers of k genomes, yielding two major groups. The first corresponds to the core genome (k = 25 genomes) and the second to the orphan genes (k = 1 genome), with 1,871, and 4,000 genes, respectively (**Fig. S2B**). Furthermore, we constructed the maximum likelihood tree from the accessory genome elements, which makes it possible to visualize the relatedness of strains based on their pan-genome composition and genes shared by different strains (**Fig. 2B**). This analysis indicates that strains belonging to groups B, C and D are also clustered together and shows that SMQ-1335 and 862_7 strains are very closely related, differing only by a few genes (**Fig. 2B**) and presenting an ANI of 99.94% (**Fig. 2A**). Both strains were isolated from cheese made in different regions (**Table S4**). Moreover, FME43 and FME45 strains, both isolated from the same cheese in this study, belong to group B with an ANI of ~98.8%, while their gene content differs by about 10%. These strains are thus closely related but their pan-genome differs significantly.

**Figure 2.**
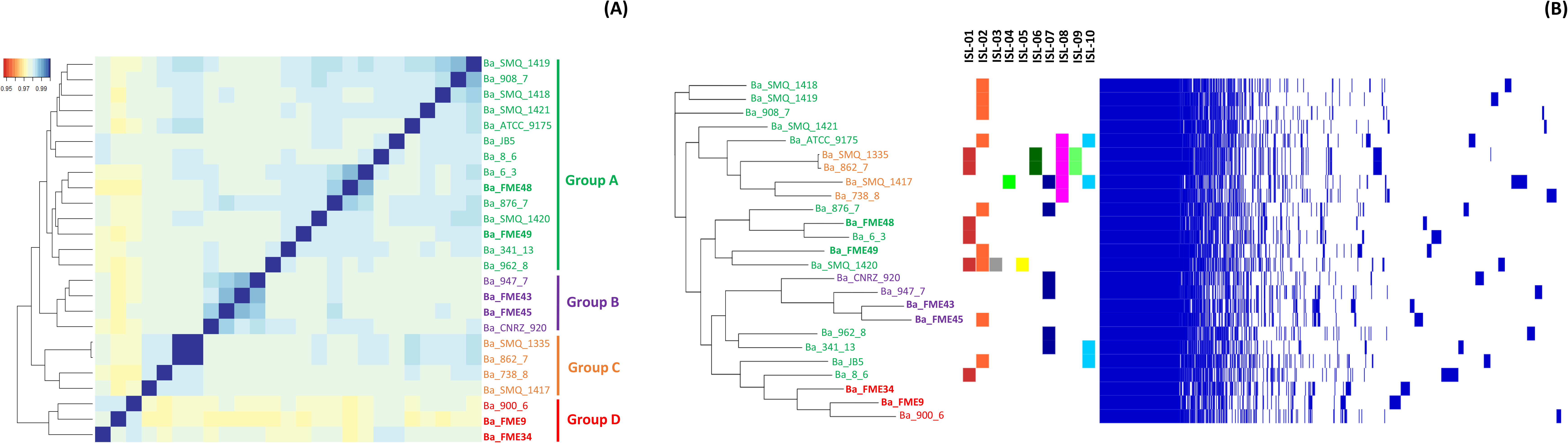
ANI and pan-genome analyses of 25 *B. aurantiacum* cheese strains. Strains from this study are highlighted in bold. **(A)** Phylogeny based on ANI values showing the presence of four groups marked in green (Group A), violet (Group B), orange (Group C) and red (Group D). **(B)** Pan-genome analysis. Left panel: maximum likelihood tree constructed from the accessory genome elements; middle panel: distribution of several horizontal gene transfer (HGT) regions described as islands [26]; Right panel: gene presence-absence matrix showing the presence (blue) and absence (white) of orthologous gene families.

In previous studies, potential horizontal gene transfer events were proposed to have occurred between *Brevibacterium* and several Actinobacteria [26–28]. From these studies, we selected ten regions containing genes involved in different metabolic functions and studied their distribution within the 25 *B. aurantiacum* genomes available (**Fig. 2B)**. This analysis shows that, except for islands 3, 4 and 5, which are present in only one strain each, the other islands are widely spread out in the different *B. aurantiacum* strains. Concerning iron transport systems, which are carried out by islands 1 and 2, they are distributed in six and nine strains, respectively, and seem to exclude each other (**Fig. 2B**), probably to avoid the cost of their overload [29]. Eleven strains do not contain any of these additional genes, indicating that although they may confer a selective advantage, alternative systems exist.

Finally, we characterized three *S. equorum* strains, which, together with the other genomes of this species available in the NCBI database (**Table S4**), make a total of 43 strains. The tree based on ANI analysis revealed two well-separated groups of 39 and four strains, indicated as Groups I and II, respectively (**Fig. 3A**). Strains belonging to Groups I and II display an ANI > 98% within their group, but an ANI of ~95% with those of the other group. Group I, which is the largest, may be subdivided into three subgroups (A, B and C), sharing an average of 99.5% ANI intra-subgroups and differenciated by over 98.7% ANI inter-subgroups. While Group II contains only dairy strains, Group I also contains strains of cattle and several other environments (**Fig. 3A**, **Table S4)**. Pan-genome analysis of *S. equorum* indicated an open pan-genome with up to 7,000 genes (**Fig. S2C**). The analysis of the number of genes present in different numbers of k genomes revealed two major groups consisting of the core genome (k = 43 genomes) and the orphan genes (k = 1 genome), with 1,868, and 2,500 genes, respectively (**Fig. S2D**), which reflects a moderate level of genetic diversity in this species. Pan-genome clustering produced several groups, which were tentatively linked to metadata and ANI (**Fig. 3B**). First, it confirmed the distinction of Group II (FME19, White_SAM, OffWhite_SAM and BC9), whose strains mainly differ from each other by their content of mobile elements (prophages, potential plasmids, etc.), while the rest of their genome is nearly identical (2,292 genes > 99.9% identity). Furthermore, seven strains isolated in different cheeses and countries (France and the U.S.) appear to be highly related (908_10, Mu_2, 876_5, 862_5, 962_6, 947_12 and 738_7; **Fig. 3B**). They share ~2,500 almost identical proteins (> 99.7%), compared to ~1,400 with the other strains of this subgroup. Their pan-genomes mainly differ by mobile elements, including potential prophages, plasmids and the number of hypothetical proteins. Finally, while several cattle strains may also form distinct groups such as those of the ANI group B, several cheese and cattle strains appear to be related.

**Figure 3.**
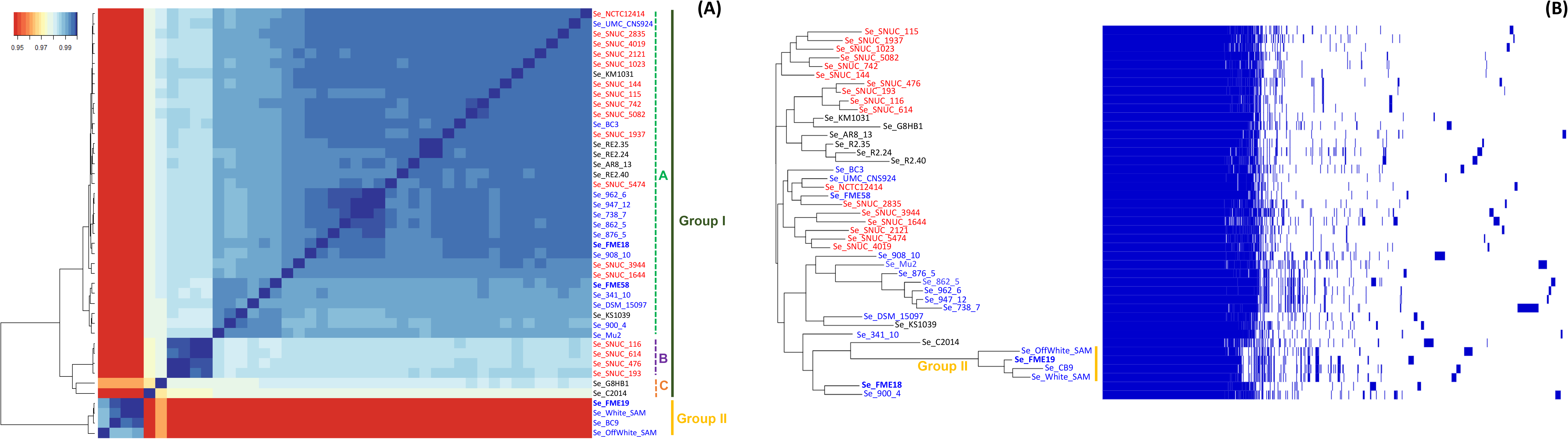
ANI and pan-genome analyses of 43 *S. equorum* strains. The origin of the strain is indicated by color: blue (dairy), red (cattle) and black (other). Strains from this study are highlighted in bold. **(A)** Phylogeny based on ANI values showing the presence of two groups potentially representing two subspecies; four subgroups in Group I (A, B, C and D) are presented by colored lines. **(B)** Pan-genome analysis. Left panel: maximum likelihood tree constructed from the accessory genome elements; right panel: gene presence-absence matrix showing the presence (blue) and absence (white) of orthologous gene families.

### Overview of halophilic species in cheese rind metagenomes

In this study, we isolated and characterized the genomes of strains belonging to 26 different species, ten of which are potentially new species. The availability of their genome sequences as references opens the possibility to detect and quantify their presence in shotgun metagenomic data. We therefore selected a set of 74 metagenomic samples corresponding to cheese rinds from different types, including 49 from this study and 25 from former studies [24, 25] (**Table S5**). In order to provide a comprehensive overview of halophilic and halotolerant species in these samples, we completed our set with 40 supplementary genomes of related species isolated from cheese in previous studies. The percentages of reads corresponding to the 66 reference genomes, which were mapped on the 74 metagenomic samples, are presented in **Table S6**. Among the 74 cheese rinds analyzed, only five have no detectable level of halophilic or halotolerant bacteria. Interestingly, more than half of the samples (42) present more than 10% of reads from these bacteria, showing their importance in cheeses.

Among Gram-positive bacteria, *Brevibacterium* and *Brachybacterium* species are widely distributed, especially in natural and washed cheese rinds (**Fig. 4**). *B aurantiacum* is the most frequently detected, in about 70% of the samples, and its amount exceeds 10% in six cheese rinds. Interestingly, the new species of *Brevibacterium* isolated here, represented by FME37, is also frequently detected in cheese rind metagenomes (half of the samples) and exceeds 5% of the reads in three cheese rinds (**Table S6**). Similarly, the potential new species represented by *Brachybacterium sp.* FME24 is detected in 12% of the samples, showing the potential relevance of this species in cheese ecosystems. Both species of *Corynebacterium, G. arilaitensis*, *A. casei* and *M. gubbeenense* are also frequent (present in more than 17% of the samples) and sometimes abundant (more than 5% of the reads) in our dataset (**Table S6**). Additionally, coagulase-negative *Staphylococci* are particularly present in several natural cheese rinds (**Fig. 4**). Among the four species, *S. equorum* is the most frequent (present in 31% of the samples) and abundant one (more than 5% of the reads in two samples). Finally, several other Gram-positive species are detected at low frequency (between 1 and 13% of the sample) and at low abundance (less than 1% of the reads) in our dataset (**Table S6**).

**Figure 4.**
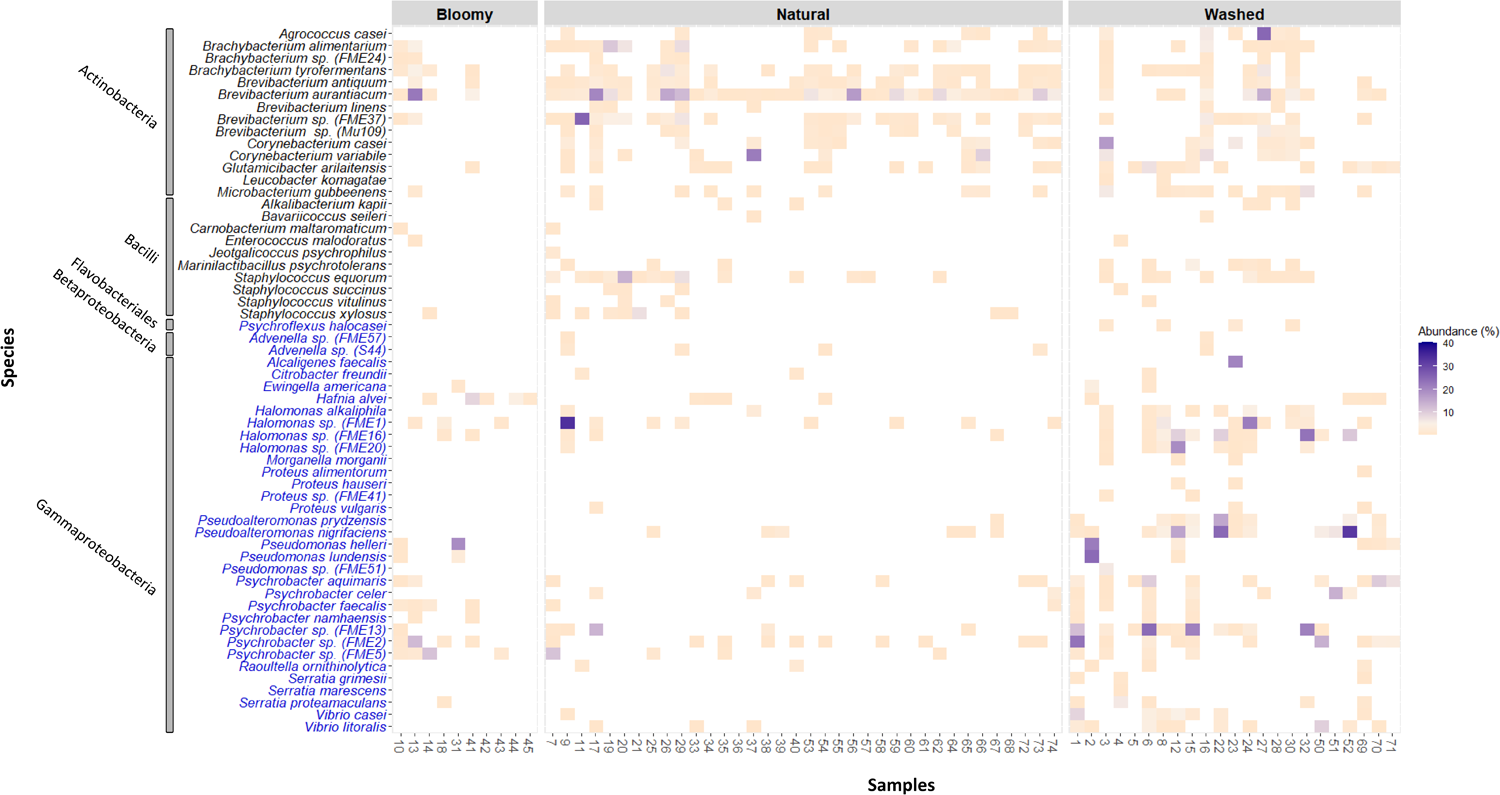
Heatmap depicting the relative abundance (%) of halophilic and halotolerant species in 74 cheeses. Samples are ordered according to rind types, as indicated by upper labels (bloomy, natural and washed). Bacterial species are ordered according to their taxonomical class and whether they belong to the Gram-positive or Gram-negative groups, by the color of their name, black and blue, respectively.

Furthermore, several Gram-negative species appear to be frequent and dominant in a significant number of samples, especially in washed cheese rinds (**Fig. 4**). In particular, different species of *Halomonas, Pseudoalteromonas* and *Psychrobacter* are detected in 13 to 32% of the samples, and several species, including new species isolated in our study, exceed 10% of the reads (**Table S6**). Three additional Gram-negative species, *V. casei*, *V. litoralis* and *H. alvei,* are also relatively frequent (11 to 18% of the samples) and sometimes abundant (more than 5% of the reads mapped). Both *Vibrio* species are mainly found in washed rinds, whereas *H. alvei* is present in bloomy rinds (**Fig. 4**). Additionally, the *Pseudomonas* FME51-like species is only detected in the cheese sample it was isolated from (Sample 3), while *P. helleri* and *P. ludensis* are detected in seven and four samples, respectively, and sometimes at high levels (up to 25% of the reads, **Table S6**). Finally, the other Gram-negative species are detected at low frequency (between 1 and 8% of the samples) and at low abundance.

### Co-occurrence relationships among bacterial species

Exploratory network and correlation analyses were performed to investigate the co-occurrence among cheese halophilic and halotolerant bacteria in order to identify combinations of species and ecosystem structuration (**Fig. 5**). Only species present > 1% in at least two cheese sample were plotted. As previously demonstrated (**Fig. 4**), Gram-positive species are more closely related in cheese with natural and washed rinds, while Gram-negative species are more closely related in cheese with washed rinds (**Fig. 5A**). Overall, the different species appear to present a higher level of co-occurrence within their group than outside (**Fig. 5A and 5B**; P value < 0.05). We observed that species belonging to the same genus are often found together, such as *Brevibacterium, Corynebacterium, Pseudoalteromonas, Halomonas, Psychrobacter, Pseudomonas* and *Vibrio* species (**Fig. 5B**). Additionally, we noted a positive correlation with *Brevibacterium* species, *Brachybacterium tyrofermentans* and *Agrococcus casei*. Finally, we highlight here the high level of co-occurrence between some species of *Psychrobacter* and *Vibrio*, as well between several species of *Halomonas* and *Pseudoalteromonas* (**Fig. 5B;** P value < 0.05).

**Figure 5.**
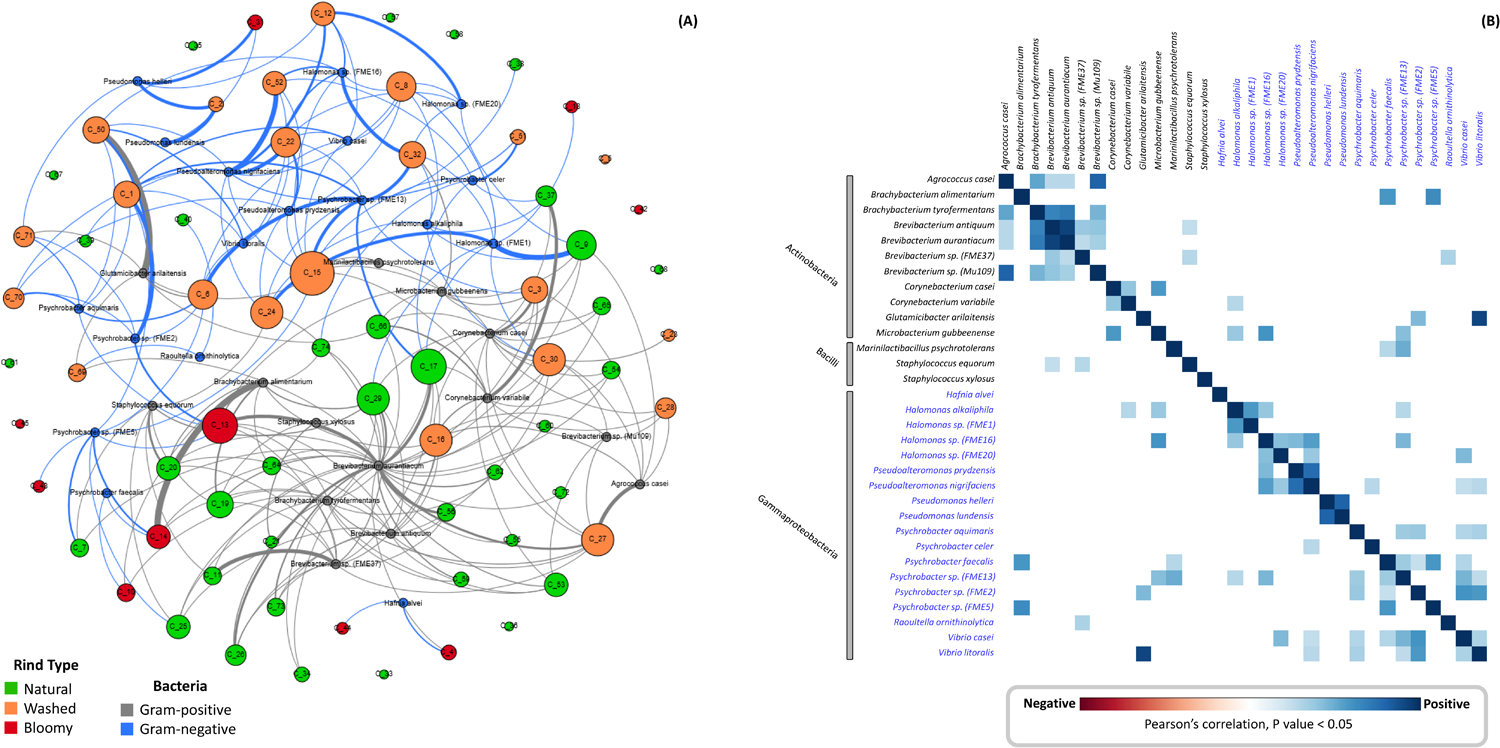
Relationships between halophilic and halotolerant bacteria detected by metagenomic analysis in cheese rinds. **(A)** Network summarizing the relationships between bacterial species and 74 cheese rind samples. Nodes represent species and cheese samples. For sample nodes, different colors (green, orange and red) are used to differentiate cheese rinds (natural, washed and bloomy, respectively). For species nodes, grey and blue are used to differentiate Gram-positive and Gram-negative strains, respectively. Connecting edges indicate the detection of given species in the samples and are colored according to whether they belong to the Gram-positive or Gram-negative group. Only edges corresponding to species detected at 1% relative abundance in the samples are shown. Sizes for cheese nodes and edges are proportional to in-degree (i.e., total occurrence of a species in the whole dataset). **(B)** Correlation matrix between halophilic and halotolerant species described in this study. Bacterial species are ordered according to their taxonomical class and whether they belong to the Gram-positive or Gram-negative groups, by the color of their name, black and blue, respectively. Only species present > 1% in at least two samples of cheese rind were plotted. Measurements are computed with Pearson’s test (P value < 0.05) and coefficient values are depicted using the following color gradient scale: red indicates negative correlations and blue indicates positive correlations.

## Discussion

We isolated halophilic and halotolerant bacteria from the rinds of 13 traditional French cheeses by selecting colonies of different morphotypes on media containing 1 to 10% salt. Total halophilic population count was evaluated on media with three different basic compositions (HM, LH and MB), usually used in the study of environmental halophiles, and the effect of increased amounts of salt was tested on MB. Counts obtained with MB and LH media were similar, whereas they were lower on HM and MB+8% NaCl (**Figure S1**), possibly due to its higher salt concentration. Moreover, incubation temperatures in the range of 20 to 30°C had no effect on global bacterial counts. In most rind samples, halophilic and halotolerant populations were 7-8 log CFU/g on the former media, which is a range similar to those reported in other studies using different culture media supplemented with salt, such as Milk Plate Count Agar (MPCA) with 5% salt [30, 31], Trypticase Soy Agar (TSA) supplemented with 4% NaCl [32] and Brain Heart Infusion (BHI) [33]. These data, together with the fact that most isolates were able to grow on all tested salt-based media, would indicate that the choice of the media is not determinant in the study of halophilic and halotolerant bacteria in cheese. Nevertheless, we did not always obtain the same species in the parallel plate isolation, suggesting that the use of different media could favor, differently and sufficiently (without being sharply selective), the growth of different populations and thus allow a greater variety of species to be isolated. From ~320 isolates, we selected 35 strains with unique 16S-*rpoB* sequences corresponding to 20 Gram-positive and 15 Gram-negative strains (**Table 2**). Their genome sequences were determined and ANI analysis showed that 12 of these isolates belong to two and eight potentially new species of Gram-positive and Gram-negative bacteria, respectively.

### Halophilic and halotolerant as food bacteria

The availabilty of the genomic sequences of halophilic and halotolerant cheese bacteria offers the opportunity to reliably detect and quantify their corresponding populations in cheese metagenomic samples at a level of 0.1% of DNA. This relative abundance level corresponds to subdominant populations, which may reach 10^5^-10^6^ CFU/g for cultivable species in several types of cheeses, a level compatible with a significant metabolic activity that could impact cheese technology.

The development of Gram-positive species belonging to Actinobacteria and Firmicutes in cheese rinds during ripening is well established and has been previously reported in different types of cheeses all over the world (**Additional File 2**). A metagenomic search showed that species like *B. tyrofermentans, B. aurantiacum, Corynebacterium* species and *G. airilaitensis* were detected in 25 to 70% of samples at maximal levels of 6 to 45% of the reads (**Table S6**), confirming their wide distribution in cheese rinds and their potential importance in cheese technology. Interestingly, we isolated a new species of *Brevibacterium* (FME37 strain) that is widely distributed in metagenomic samples and sometimes detected at high levels (> 20%), suggesting a role in cheese.

*B. aurantiacum* is the most frequent and abundant Actinobacteria found in our metagenomic samples. This frequency was probably boosted by the over 40-year-old history of the use of this bacteria as an adjunct to take advantage of its various technological properties [34, 35], although the persistense of adjuncted strains is questioned [36, 37]. This species was subject to the most detailed genomic analysis, and 19 cheese genome sequences were available at the time of this study. The comparison of these genomes, in addition to the six provided here, confirmed the substantial genetic diversity within *B. aurantiacum* and its plasticity, which might be reflected in the diversity of color, aroma, lipolytic and other technological factors described for this bacteria. Finally, the study of the different genomic islands - characterized earlier – suggests that their roles are not crucial for the development of this species in cheese since a significant number of strains are not concerned by these additional factors. In particular, from the two strains FME43 and FME45 (isolated from the same cheese), only one contains iron acquisition genes on ISL2 (**Fig. 2B**), which were described as being important to develop within the cheese surface habitat [27, 38]. The alternative distribution of the two different iron acquisition systems on ISL1 and ISL2 in about half of the strains may suggest that their presence is also a metabolic load, whereas alternative strategies probably exist.

Furthermore, we isolated three coagulase-negative staphylococci species, a type of bacteria commonly isolated from cheese. In agreement with a previous metagenomic study [39], we found that *S. equorum* is the most frequent and abundant species, while *S. succinus* and *S. vitulinus* are more scarce (**Fig. 4**). *S. equorum*, which is also used as an adjunct in cheese to improve its texture and contribute particular flavors (**Additional File 2**), has been extensively studied for safety reasons, and 40 genomes from food, cattle and clinical samples were available at the time of this study. Their ANI analyses, together with our three isolates, suggest that *S. equorum* could be subdivided into two subspecies (**Figure 3A**), the second one being represented by four cheese strains. The combined ANI and pan-genome analyses of these strains (**Fig. 3A and 3B**) indicates that they mainly differ by their mobile element content (prophages and potential plasmids; **Fig. 3B**), whereas the rest of their genomes are nearly identical, suggesting a recent common origin for their use in cheese. Further investigations will be required to determine if the two potential subspecies express different technological properties, including phage resistance. In the major group of strains (Group IA), many isolates from cheese could not be clearly differenciated from those from cattle, suggesting their animal origin, except a group of seven cheese strains (**Fig. 3B**). The latter ones mainly differ by their mobile elements, which could be the result of starter culture selection. However, further studies will be necessary to demonstrate that these strains followed a drift and are now specifically growing in cheese rinds.

Additionally, we isolated three other less documented Firmicutes, including *Marinilactibacillus psychrotolerans* and *Carnobacterium mobile* (occasionally found in cheese), and *Oceanobacillus oncorhynchi*, a halotolerant bacteria sometimes isolated from Asian salted food (**Additional File 2**). Whereas *M. psychrotolerans* was detected in around 13% of our metagenomic data with a maximum of 4% of the reads, *C. mobile* and *O. oncorhynchi* were not detected, including the samples from which they were isolated (**Table S6**). These results indicate that the size of the population of these two species was very small and that they may not play a significant role in cheese technology.

In addition to these halotolerant Gram-positive bacteria, we obtained several species of Gram-negative bacteria that have not yet aroused keen interest in technological developments. Remarkably, Gammaproteobacteria from the genera *Halomonas, Pseudoalteromonas* and *Psychrobacter* were the most frequently isolated (representing 60% of the Gram-negative isolates obtained in this study), and they often represent high relative abundance in washed cheese rinds (**Fig. 4**). This observation is in agreement with recent culture-independent analyses [24, 25], and their presence in cheeses of good quality supports the possibility that they may have a positive impact during ripening, while several authors consider them as contaminants (**Additional File 2**). Nevertheless, the characterization of the potential role of these genera in cheese technology remains to be explored. Finally, we isolated species of *Pseudomonas*, *Proteus*, *Hafnia, Vibrio* and *Advenella*. All these genera have already been reported in cheese rind communities and the abundance of several of these species may support further interest for their role in cheese ripening (**Additional File 2**). Currently, *Hafnia alvei* is the only Gram-negative bacterium used as a ripening starter in the cheese-making process and it was detected here at level of 4.5 and 8.5% in two bloomy cheese rinds (**Table S6**), where it was probably added as an adjunct culture.

### New insights into cheese microbial ecology

Interstingly, the development of Gram-negative bacteria appears to be greater in soft cheese, and also favored by the washing and smearing processes of the rind, which correspond to cheeses displaying higher moisture [40], as roughly depicted in **Figure 1**.

Remarkably, the present metagenomic analysis discriminates populations at the level of species, which is not always possible with amplicon sequencing that is limited by the lack of significant divergence, especially for Gram-negative species such as those presented in this study. Consequently, the present set of data uncovers the world of halophilic and halotolerant bacteria in cheese rinds with a high level of resolution, and reveals, in particular, the co-occurence of a number of species, including closely related species hardly distinguishable by marker gene analysis. For example, species of *Brevibacterium, Corynebacterium, Halomonas, Psychrobacter, Pseudoalteromonas* and *Vibrio* are found co-occuring with at least another species of the same genus (**Fig. 5B**). The coexistence of such a variety of species reflects the fact that these ecosystems are open to the microbial environment, which is likely to be resilient to their production conditions and develop with salt as a key driver. The presence of related species, which are thus likely to carry out similar metabolic functions, will lead to a functional redundancy, a factor that was proposed to be of primary importance in the the resilience of ecosystems submitted to changes or pressures [41, 42]. In the context of traditional productions, changes could encompass modification of milk quality, technological issues and phage attacks, thus structuring the ecosystem and maintaining a variety of microorganisms. However, while the presence of diverse microorganisms in the processing environment may increase the ecosystem’s capacity to respond to changes, it may also modify the organoleptic properties of cheeses [43]. Further studies will be required to understand the interactions that occur between these microorganisms and their role in the development of cheese, as well as the development of the rich and various organoleptic properties of traditional products.

The present study made it possible to isolate 26 different species, ten of which belong to still undescribed species, although they are frequent and abundant in various cheeses. These strains could be used as references to promote advances in functional studies of this particular world driven by salt addition and to jointly contribute to safe and efficient cheese development.

## Materials and methods

### Cheese sampling

A total of 13 cheese samples were selected in this study (**Table 1**). The cheeses were purchased from a local supermarket and their rinds were sampled in portions (1 g) with a sterile knife and frozen at −20°C until further analysis.

### Enumeration and isolation of halophilic bacteria from cheeses

To enumerate and isolate halophilic strains, we used Marine Broth (MB; Difco, Sparks, USA), Long and Hammer Agar (LH; [44]) and *Halomonas* Medium (HM; [45]). Different concentrations of salt were supplemented in MB (0, 4, 6 and 8% NaCl). In order to prevent fungal growth, Amphotericin B (Sigma-Aldrich, St. Louis, MO, USA) was added to a final concentration of 20 μg/ml (50 mg/ml stock solution of Alfa Aesar ™ Amphotericin B from *Streptomyces nodosus* in DMSO). Cultivable bacterial strains were enumerated using serial dilutions of homogenized cheese samples in sterile 0.9% NaCl solution. Population counts of cheese rinds were determined by 10^−3^ to 10^−7^ dilutions and incubated 48-72 h at 20, 25 and 30°C. For each cheese, the plates with a bacterial count comprised between 20 and 200 clones were selected for isolate characterization. An initial selection of apparently different isolates (morphotypes) was performed based on colony morphology (color, shape, elevation, pigmentation and opacity). A representative of each morphotype was then restreaked on a new plate for subsequent DNA extraction.

### Identification of isolated morphotypes

The selected clones were collected with a sterile loop and mixed in a tube containing 300 μl biomol water, 100 mg of 0.1 mm-diameter zirconium beads and 100 mg of 0.5 mm-diameter (Sigma, St. Louis, MO, USA). The tube was then vigorously shaken in a bead-beater (FastPrep-24, MP Biomedicals Europe, Illkirch, France) for 20 s at 4.5 m/s. The supernatant of this lysis was used directly for DNA amplification. The species assignation was performed by sequencing the 16S rRNA and the *rpoB* genes. The 16S rRNA gene was amplified using 27-F (5′-AGAGTTTGATCATGGCTCA-3′) and 1492-R (5′-TACGGTTACCTTGTTACGACTT-3′) [46]. The *rpoB* gene was amplified using primers VIC4 (5′-GGCGAAATGGCDGARAACCA-3′) and VIC6 (5′-GARTCYTCGAAGTGGTAACC-3′) [24, 47].

Thermal cycling conditions applied for both were (i) 1 min at 94°C to initial denaturation; (ii) 30 cycles of 1 min at 94°C to denaturation, 0.5 min at 56°C to primer annealing, 1.5 min at 72°C to initial elongation; and (iii) 5 min at 72°C to final elongation. DNA amplifications were separated on 0.8% agarose gel. The PCR products were purified using the ExoSAP-IT (Thermo Fisher Scientific, Waltham, MA, USA) and sent for sequencing to the service provider (Eurofins Genomics, Ebersberg, Germany). Once received, the sequences were analyzed via the NCBI BLAST tool [48] to obtain a taxonomic classification for each isolate.

### Genomic DNA extraction and sequecing

For each unique isolate, after cultivation in MB for 48 h at 25°C, DNA was extracted from the bacterial cells according to the protocol described by Almeida *et al.* [24] with some modifications. Briefly, we used an enzymatic lysis step followed by protein precipitation by adding potassium acetate. DNA was precipitated at −20°C after the addition of 0.1 volume of 3 M sodium acetate and two volumes of cold absolute ethanol to the upper phase. After centrifugation (30 min at 12,000 *g* and 4°C), the DNA was dried in a laboratory hood and resuspended in TE 1X buffer. The DNA concentration and quality was evaluated using a NanoDrop ND-1000 spectrophotometer (NanoDrop Technology Inc., Wilmington, DE, USA). Additionally, 5 μL of DNA were loaded on 0.8% agarose gel and visualized after migration by ethidium bromide staining.

DNA sequencing was carried out on an Illumina HiSeq at GATC-Biotech (Konstanz, Germany) in order to generate paired-end reads (150 bases in length). For each strain, the paired-end reads were merged and *de novo* assembly was performed using SPAdes, version 3.9 [49]. Only contigs with length◻>◻300 bp and coverage >100 were considered for further study. Annotations were performed using the Rapid Annotation using Subsystem Technology server [50].

### Phylogenetic analysis

For species assignation, evolutionary trees were built using the 16S rRNA and *rpoB* genes. To delineate species, we used a threshold of over 99% identity for 16S rRNA genes with type or well-defined strains [51], and of above 97.7% identity for *rpoB* nucleotide sequences [52]. Further phylogenetic analyses were performed using ClustalX 2.1 [53] and MEGA7 [54]. The trees were built using the Neighbor-Joining method [55] with 1,000 bootstrap replicates [56]. Lastly, using genomic sequences, we determined the Average Nucleotide Identity (ANI) using JSpeciesWS [57] to confirm speciation of the different isolates. For new species delineation, we used the recommended cut-off point of 95% ANI [58].

### Pan-genome of *Brevibacterium aurantiacum* and *Staphylococcus equorum*

ANI and pan-genome analyses were estimated for 25 genomes of *Brevibacterium aurantiacum* and 43 genomes of *Staphylococcus equorum* (**Table S4**). The ANI was performed using the ANIm method described by Richter *et al.* [59] and implemented in the Python module PYANI (version 0.2.6) (https://github.com/widdowquinn/pyani). The pan-genomes of both species were performed with Roary software (version 3.11.2; [60]) and the gene-based genome-wide association using Scoary [61]. Interactive visualization of genome phylogenies was done with Phandango (version 1.3.0; [62]).

### Cheese DNA extraction and sequencing

From the 13 samples used to isolate halophilic and halotolerant bacteria, ten were selected to analyze their total DNA. The DNAs were prepared from cell pellets obtained from each cheese sample, following a method that combines enzymatic and mechanical treatments for cell lysis and treatment with phenol/chroloroform/isoalmyl alcohol to extract and purify DNA, as previously described by Almeida *et al.* [24]. The ten DNA samples from cheese were sequenced using Illumina HiSeq2500 technology at GATC-Biotech (Konstanz, Germany), which yielded between six and eight million paired-end reads of 150-nucleotide length. Moreover, 39 additional samples from different types of cheese were sequenced using SOLiD technology, which yielded between 11-19 million single reads of 50-nucleotide length. The raw read data for all samples are available under the accession numbers listed in **Table S5**.

### Quantification of species in metagenomic samples

First, species present in each metagenomic sample were identified with the Food-Microbiome Transfert tool, an in-house designed tool managing the following tasks. Each of the 66 reference genomes (one genome per species) was mapped on the metagenomic samples with Bowtie [63] (adapted to SOLiD data, parameters were adapted to take into account intra-species polymorphisms and choose at most one mapping position per read: first 35 nucleotides mapped; 3 mismatches allowed; --all --best --strata -M 1). In order to discard reads that could have been aligned on conserved regions on a more distant genome (same genus for example) or repeated regions, BEDtools [64] and SAMtools [65] were used to filter reads and compute genome coverage. Reads mapping genomic regions that were less informative and/or that could have been acquired by gene transfer (intergenic regions, tRNA, rRNA, genes annotated as “transposase”, “integrase”, “IS”, “phage/prophage” or “plasmids”) were not taken into account. In order to select genomes whose species is present in the sample, we selected genomes with at least 50% of their genes covered by at least one read. Food-Microbiome Transfert tool was used via a web interface developed via the Python Django framework as well as web technologies such as HTML and JavaScript. Genome, metagenome and analysis data are stored on a PostgreSQL relational database. Computations were performed on the Migale platform’s calculation cluster via the Bioblend API and the Galaxy portal.

Then, to determine the abundance of the different halophilic and halotolerant species, metagenomic reads were mapped on a database containing all 66 reference genomes with Bowtie (same parameters). We selected only reads mapping on genomes selected at the previous step. Quantification was done by counting the number of read for each genome. In order to obtain comparable results between metagenomes, we downsized the samples to 5 million reads. The metadata of metagenomic samples are presented in **Table S5**.

### Statistical analysis of co-occurrence relationships

The relationship between the halophilic species was examined by performing a correlation matrix using Pearson’s test. The function ‘rcorr’ (in ‘Hmisc’ package) was used to compute the significance levels (P<0.05) and the graph were plotted using the ‘corrplot’ package for R. Only species with a relative abundance ≥1% were used to generate matrix and network correlations. Bacterial networks were explored and visualized using Gephi software 0.9.2 [66].

### Data availability

Raw genomic reads were deposited to the Europenan Nucleotide Archive under the project accesion number PRJNA501839, while Illumina and SoliD metagenomic reads under PRJNA642396 and PRJEB39332, respectively.

## Acknowledgments

This work was co-funded by the French Dairy Interbranch Organization (CNIEL, Centre National Interprofessionnel de l’Economie Laitière), Paris, France. CIK’s grant was supported by “Conselho Nacional de Desenvolvimento Científico e Tecnológico (CNPq)” – Brazil [grant number 202444/2017-1].

The authors thank all the members of CNIEL’s committee on dairy microbiology, Françoise Irlinger, Christophe Monnet, Céline Delbès and Delphine Passerini for stimulating discussions. The authors are grateful to Mathieu Almeida and Nicolas Pons for their help with SoliD data management and Thibaut Guirimand, Quentin Cavaillé and Charlie Pauvert for their involvement in the Food-Microbiome Transfert tool.

## Authors contributions

PR conceived the study and its experimental design. BD and BFK collected samples, isolated and identified strains. CIK performed genomic and metagenomic analysis. AB contributed to genomic analysis. The Food Microbiome team supported overall cheese rind metagenomic analysis. CIK and PR analyzed the data and wrote the manuscript. PR supervised the project.

## Supplementary Material

**Additional File 1**. This supplementary information documents the identification of halophilic and halotolerant isolates using the phylogenetic analysis of 16S rRNA and *rpoB* genes.

**Additional File 2**. This supplementary information documents the main bacterial species of halophilic and halotolerant bacteria in food and their potential roles.

**Figure S1.** Total viable counts in different salt media and temperatures in log CFU/g from 13 cheese rind samples.

**Figure S2**. Pan-genome analyses **(A)** Accumulated number of new genes in the pan-genome and genes of *B. aurantiacum* attributed to the core-genome are plotted against the number of added genomes. **(B)** Accumulated number of genes in k genomes are plotted against the number of k genomes. The number of core genes present in all 25 *B. aurantiacum* genomes can be observed with k = 25 genomes. **(C)** Accumulated number of new genes in the pan-genome and genes of *S. equorum* attributed to the core-genome are plotted against the number of added genomes. **(D)** Accumulated number of genes in k genomes are plotted against the number of k genomes. The number of core genes present in all 43 *S. equorum* genomes can be observed with k = 43 genomes.

**Table S1.** Potential assignment of 35 representative isolates using 16S rRNA and *rpoB* genes.

**Table S2**. General features of the 35 genomes sequenced in this study.

**Table S3**. Average Nucleotide Identity (ANI) with close relatives of strains that marker genes (16S rRNA and *rpoB*) were unable to reliably assign to a described species. ANI values greater than or equal to 95% are shown in green.

**Table S4**. Metadata of 25 *Brevibacterium aurantiacum* and 43 *Staphylococcus equorum* used for pan-genome analyses.

**Table S5.** Metadata of 74 cheese rind metagenomes used to map the halophilic and halotolerant genomes selected in this study.

**Table S6.** Abundance (%) of each halophilic and halotolerant species in 74 cheese rinds. The table includes the abundances corresponding to the taxonomical order of the species, the type of bacteria (Gram-positive or Gram-negative) and the total halophilic and halotolerants in each sample. Corresponding statistics on the number of samples containing more than 0.1, 0.5, 1, 5 and 10% are also available.

