## Supplementary material for "Unraveling the world of halophilic and halotolerant bacteria in cheese by combining cultural, genomic and metagenomic approaches": Figure S1

### Slide 1
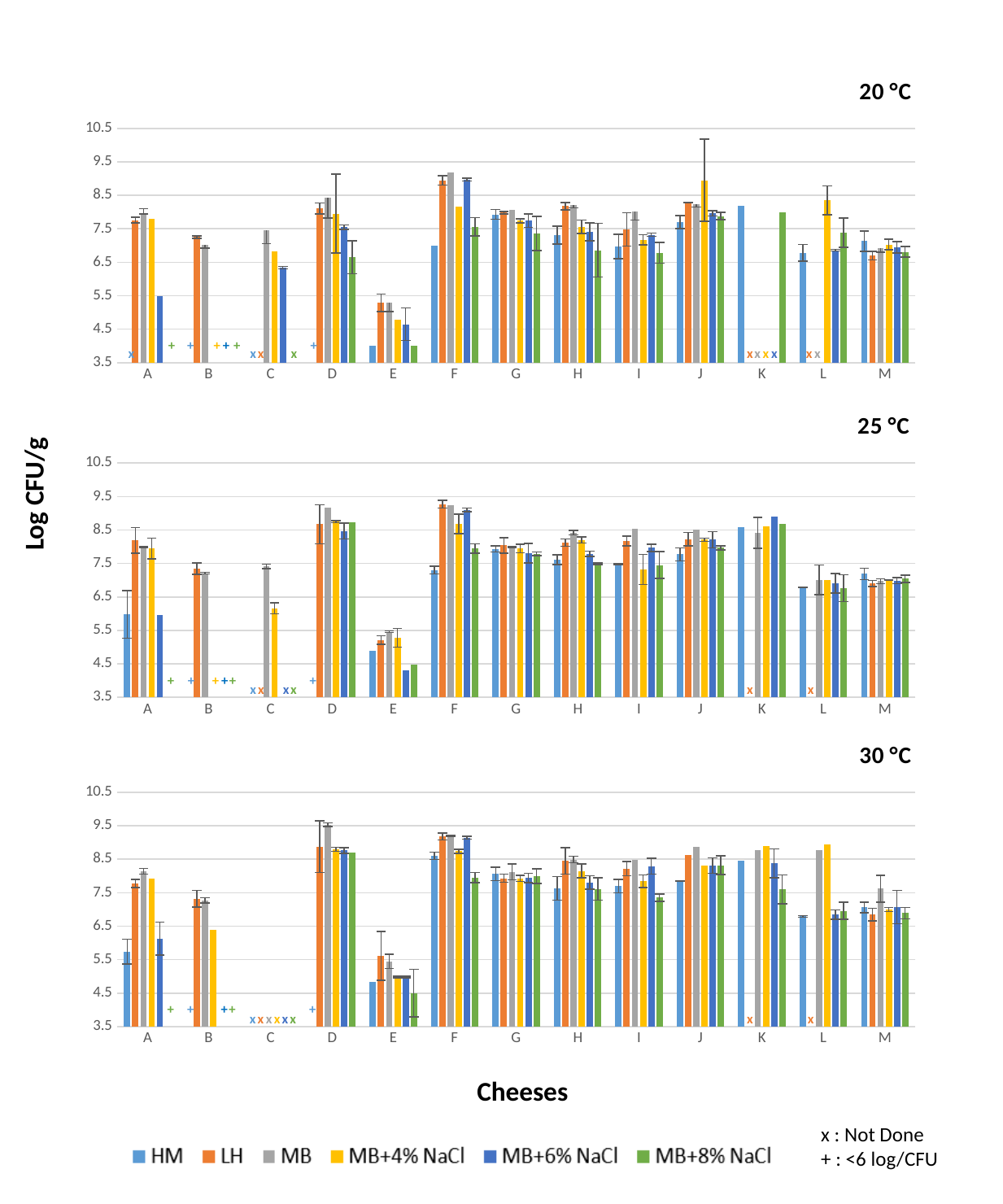

#### Chart: 20 °C
| Category | HM | LH | MB | MB+4% NaCl | MB+6% NaCl | MB+8% NaCl |
|---|---|---|---|---|---|---|
| A | 0.0 | 7.76 | 7.97 | 7.79 | 5.48 | 0.0 |
| B | 0.0 | 7.25 | 7.03 | 0.0 | 0.0 | 0.0 |
| C | 0.0 | 0.0 | 7.45 | 6.83 | 6.34 | 0.0 |
| D | 0.0 | 8.11 | 8.42 | 7.95 | 7.55 | 6.65 |
| E | 4.0 | 5.29 | 5.29 | 4.78 | 4.65 | 4.0 |
| F | 7.0 | 8.94 | 9.18 | 8.17 | 8.97 | 7.56 |
| G | 7.93 | 7.98 | 8.07 | 7.74 | 7.74 | 7.36 |
| H | 7.31 | 8.18 | 8.2 | 7.56 | 7.41 | 6.86 |
| I | 6.97 | 7.48 | 8.02 | 7.17 | 7.32 | 6.78 |
| J | 7.7 | 8.29 | 8.23 | 8.95 | 7.96 | 7.88 |
| K | 8.18 | 0.0 | 0.0 | 0.0 | 0.0 | 8.0 |
| L | 6.78 | 0.0 | 0.0 | 8.35 | 6.86 | 7.38 |
| M | 7.13 | 6.7 | 6.93 | 7.03 | 6.94 | 6.81 |+
+
+
+
+
+
x
x
x
x
x
x
x
x
x
x
#### Chart: 25 °C
| Category | HM | LH | MB | MB+4% NaCl | MB+6% NaCl | MB+8% NaCl |
|---|---|---|---|---|---|---|
| A | 5.98 | 8.19 | 7.99 | 7.95 | 5.95 | 0.0 |
| B | 0.0 | 7.35 | 7.21 | 0.0 | 0.0 | 0.0 |
| C | 0.0 | 0.0 | 7.41 | 6.16 | 0.0 | 0.0 |
| D | 0.0 | 8.67 | 9.18 | 8.75 | 8.47 | 8.73 |
| E | 4.9 | 5.21 | 5.46 | 5.28 | 4.3 | 4.47 |
| F | 7.3 | 9.27 | 9.23 | 8.68 | 9.1 | 7.95 |
| G | 7.93 | 8.04 | 7.99 | 7.95 | 7.81 | 7.79 |
| H | 7.61 | 8.12 | 8.42 | 8.2 | 7.79 | 7.49 |
| I | 7.48 | 8.17 | 8.54 | 7.32 | 7.97 | 7.45 |
| J | 7.77 | 8.23 | 8.5 | 8.21 | 8.21 | 7.97 |
| K | 8.59 | 0.0 | 8.41 | 8.6 | 8.91 | 8.68 |
| L | 6.78 | 0.0 | 7.01 | 7.0 | 6.91 | 6.76 |
| M | 7.19 | 6.9 | 6.97 | 7.0 | 6.98 | 7.04 |Log CFU/g
+
+
+
+
+
+
x
x
x
x
x
x
#### Chart: 30 °C
| Category | HM | LH | MB | MB+4% NaCl | MB+6% NaCl | MB+8% NaCl |
|---|---|---|---|---|---|---|
| A | 5.74 | 7.78 | 8.14 | 7.92 | 6.13 | 0.0 |
| B | 0.0 | 7.32 | 7.27 | 6.4 | 0.0 | 0.0 |
| C | 0.0 | 0.0 | 0.0 | 0.0 | 0.0 | 0.0 |
| D | 0.0 | 8.87 | 9.53 | 8.8 | 8.76 | 8.7 |
| E | 4.84 | 5.61 | 5.45 | 4.98 | 4.98 | 4.5 |
| F | 8.6 | 9.18 | 9.2 | 8.74 | 9.14 | 7.95 |
| G | 8.06 | 7.93 | 8.12 | 7.93 | 7.94 | 7.99 |
| H | 7.63 | 8.45 | 8.49 | 8.15 | 7.81 | 7.61 |
| I | 7.7 | 8.22 | 8.48 | 7.84 | 8.29 | 7.35 |
| J | 7.85 | 8.62 | 8.87 | 8.32 | 8.31 | 8.32 |
| K | 8.45 | 0.0 | 8.78 | 8.9 | 8.38 | 7.6 |
| L | 6.79 | 0.0 | 8.76 | 8.94 | 6.85 | 6.96 |
| M | 7.06 | 6.84 | 7.62 | 7.0 | 7.07 | 6.89 |+
+
+
+
+
x
x
x
x
x
x
x
x
Cheeses
x : Not Done
+ : <6 log/CFU
