## Supplementary figures and images for "Unraveling the world of halophilic and halotolerant bacteria in cheese by combining cultural, genomic and metagenomic approaches"

### Figure S2

#### Slide 1
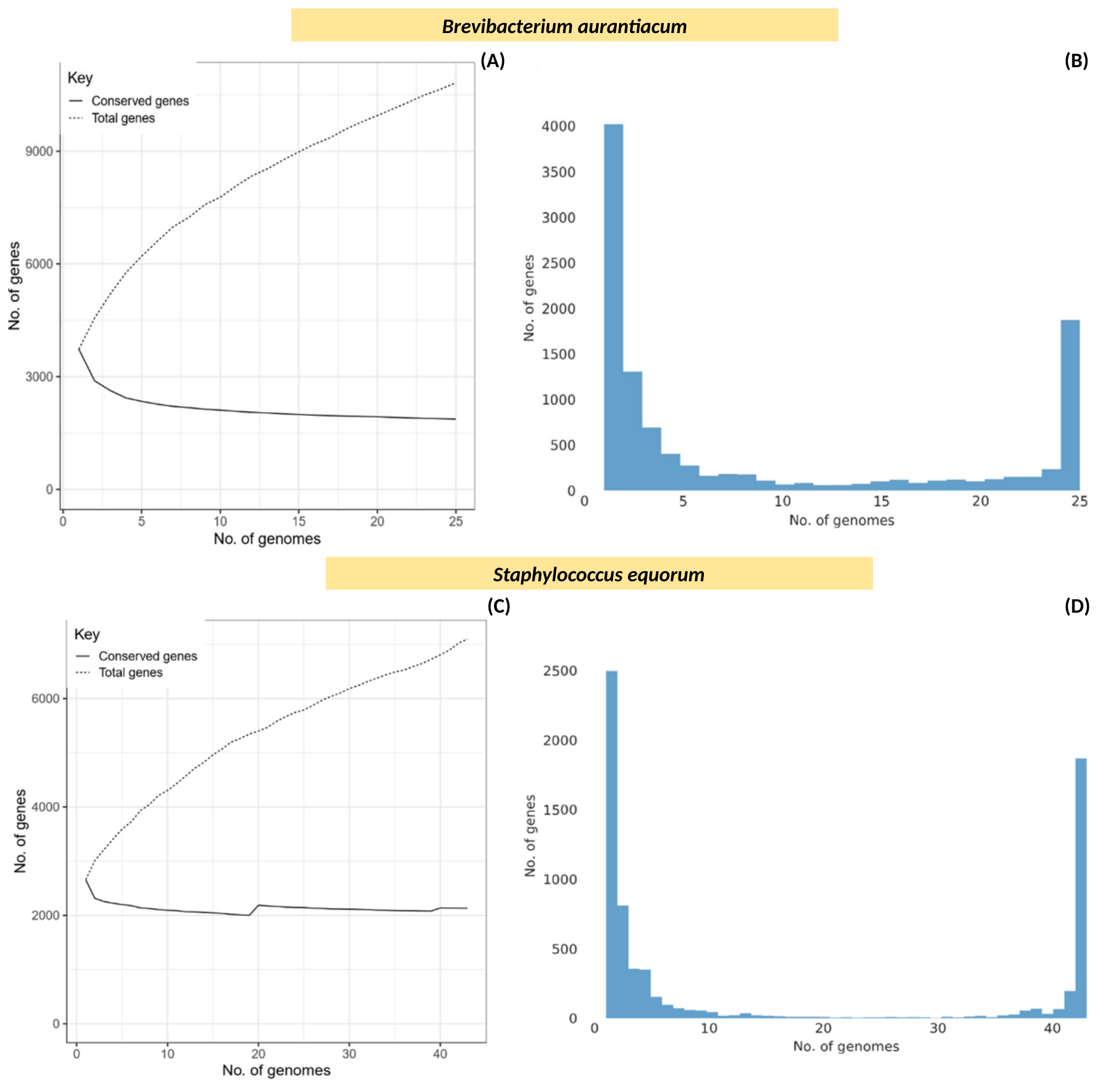

Brevibacterium aurantiacum
(A)
(B)
Staphylococcus equorum
(C)
(D)
