## Additional File 1 for "Unraveling the world of halophilic and halotolerant bacteria in cheese by combining cultural, genomic and metagenomic approaches"

**Additional File 1. Identification of halophilic and halotolerant isolates using phylogenetic analysis of 16S rRNA and *rpoB* genes.**

Among the 20 Gram-positive bacteria isolated here, twelve belongs to Actinobacteria and eight to Bacilli class (**Table S1**). The Actinobacteria were distributed in four genera: *Brachybacterium* (2), *Brevibacterium* (8), *Corynebacterium* (1) and *Glutamicibacter* (1). Among these strains, nine could be assigned unambiguously at the level of species to yield *Brachybacterium tyrofermentans* (1), *Brevibacterium aurantiacum* (6), *Corynebacterium casei* (1) and *Glutamicibacter* *arilaitensis* (1). However, the second *Brachybacterium* isolated in this study (FME24) could not be assigned reliably to a species althought its 16S rRNA share around 99% identity with the to *B. tyrofermentans* type strain, since its *rpoB* nucleotide sequence only shares 93.43% with this strain (**Table S1, Fig. A**).

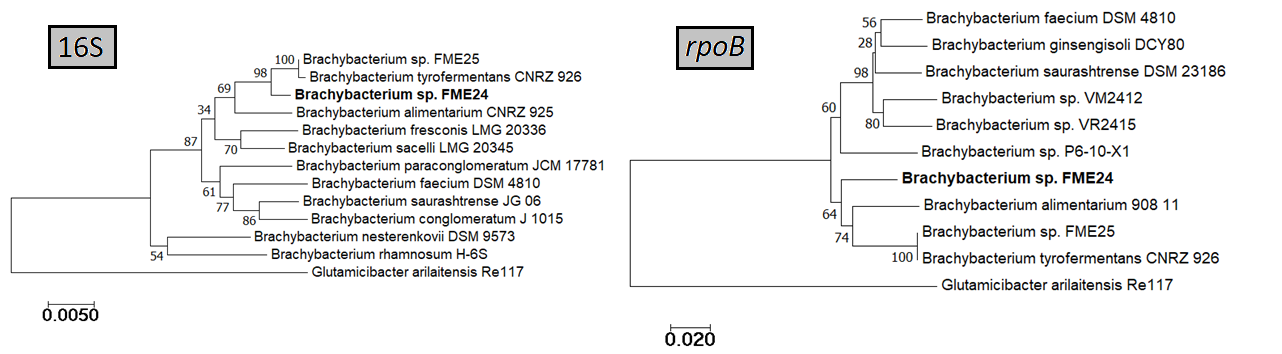

**Figure A.** Phylogenetic tree of *Brachybacterium* genus based on 16S rRNA and *rpoB* gene alignments, including FME24 and FME25 strains (isolated in this study) and their closest relatives.

Beside, the two last *Brevibacterium* (FME17 and FME37) 16S rRNA displayed only ~98% identity with the type strain of *B. antiquum,* a level which is insufficient to assign them reliably at this species. Moreover, their assignation to this species could not be confirmed by the analysis of their *rpoB* sequences, which share less than 96% identity (**Table S1, Figure B**)*.*

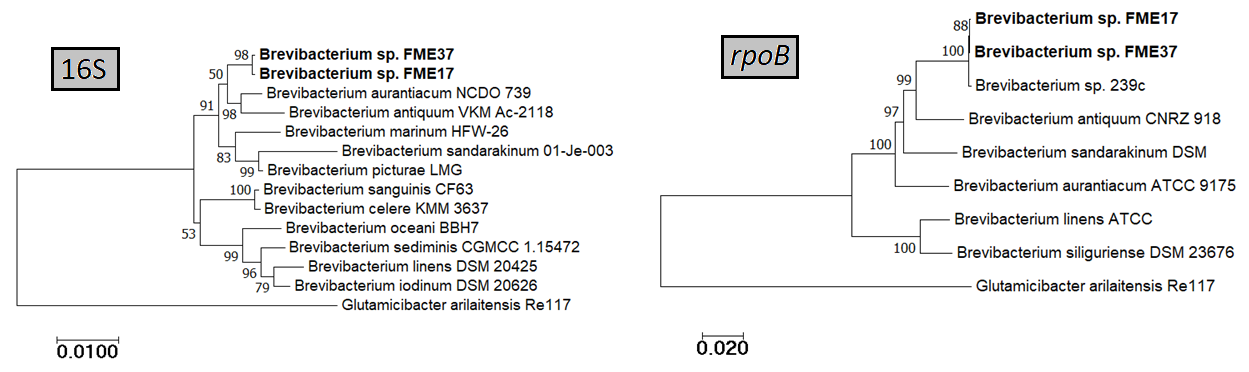

**Figure B.** Phylogenetic tree of *Brevibacterium* genus based on 16S rRNA and *rpoB* gene alignments, including FME17 and FME37 strains (isolated in this study) and their closest relatives.

Finally, isolates belonging to Bacilli class were distributed in four genera that could be unambiguously assigned at the level of species as follows, *Carnobacterium* *mobile* (1), *Marinilactibacillus psychrotolerans* (1), *Oceanobacillus orcorhynchi* (1), *Staphylococcus* *equorum* (3), *Staphylococcus succinus* (1) and *Staphylococcus vitulinus* (1) (**Table S1**).

Among the 15 strains from the Gram-negative group, one belongs to Betaproteobacteria and 14 to Gammaproteobacteria class. The Betaproteobacteria isolated in this study (FME57), possess high level of identity with the 16S rRNA of *Advenella kashmirensis*, *A. incenata* and *A. mimigardefordensis* type strains (respectively 98.9, 99.25 and 99.48%) as presented **Figure C**.

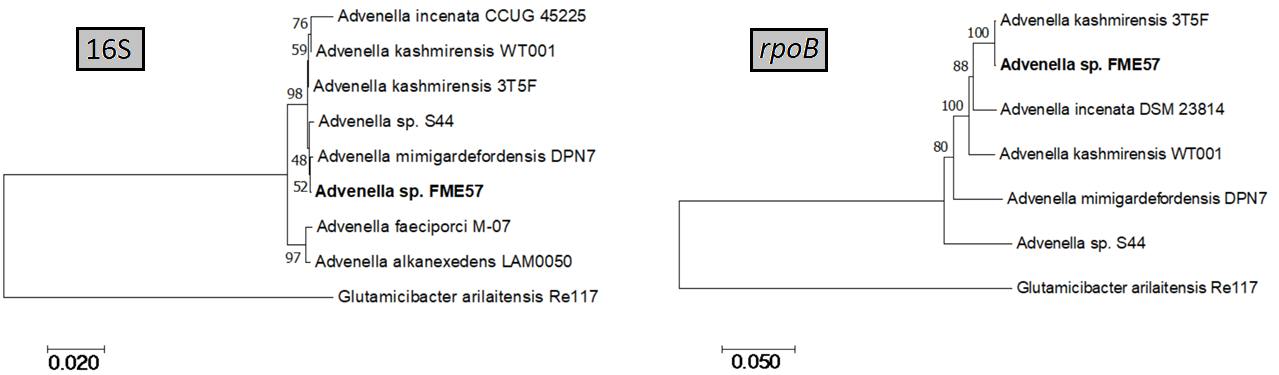

**Figure C.** Phylogenetic tree of *Advenella* genus based on 16S rRNA and *rpoB* gene alignments, including FME57 strain (isolated in this study) and its closest relatives.

Remarkably, the 16S rRNA of FME57 strain is 99.92% identical to that of *Advenella kashmirensis* 3T5F and 99.74% from *Advenella* *sp.* S44, which were isolated from cheese. Furthermore, its *rpoB* gene sequence best hit at NCBI was with *Advenella incenata* DSM 23814^T^ (96.5%) (**Table S1**), and it was only 96.26% with the *Advenella kashmirensis* WT001^T^, while still is 100% identical with that of *Advenella kashmirensis* 3T5F. These data indicated that 3T5F and FME57 strains belongs to the same species, but may not be assigned reliably to *A. kashmirensis* (**Figure C**). Concerning isolates belonging to the Gammaproteobacteria group, they were distributed in seven genera: *Hafnia* (1), *Halomonas* (3), *Proteus* (1), *Pseudoalteromonas* (2), *Psychrobacter* (4) and *Vibrio* (1). Among these strains, three could be assigned unambiguously at the level of species considering the two marker genes data to yield *Hafnia alvei* (1), *Pseudomonas lundensis* (1) and *Vibrio casei* (1) (**Table 2**). In the case of *Halomonas* FME20 strain, its 16S rRNA and *rpoB* gene sequences display high homology, respectively 99.5 and 98.5%, with those of *Halomonas zhanjiangensis* DSM 21076^T^ (**Table 2**). FME20 may thus belong to this species or be closely related to, and this hypothesis deserve further analysis to be confirmed. The two last *Halomonas* strains, FME1 and FME16, remain unassigned to the species level since their 16S rRNA and *rpoB* genes display levels of identity below the thresholds to all the defined species; however, they are closely related to JB37 and JB380 strains, respectively, which were isolated from cheese (**Table S1, Fig. D**).

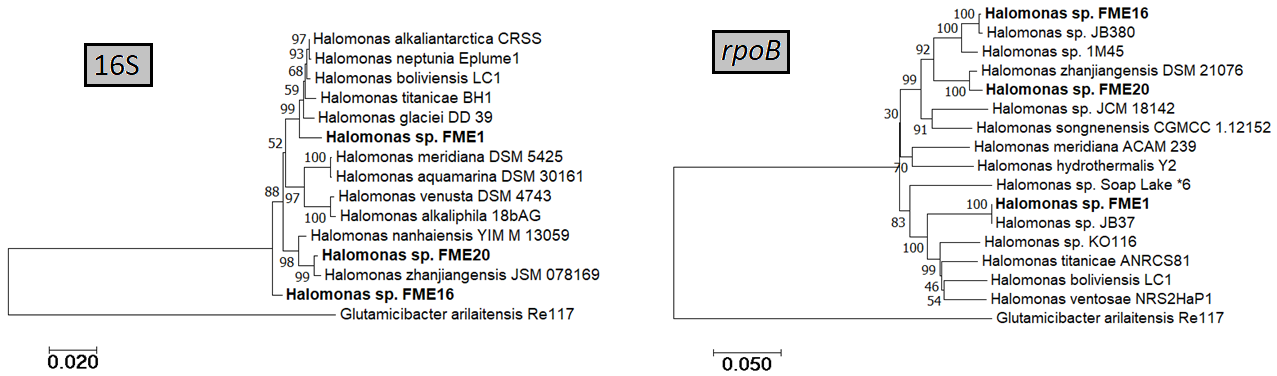

**Figure D.** Phylogenetic tree of *Halomonas* genus based on 16S rRNA and *rpoB* gene alignments, including FME1, FME16 and FME20 strains (isolated in this study) and their closest relatives.

Beside this, the *Proteus* isolate FME41 also remained unassigned because, althought its 16S rRNA shares around 99% identity with the one of type strain, *Proteus hauseri* DSM 21076^T^ , the *rpoB* nucleotide sequences of these two strains share only 93.92% (**Table 2, Figure E**).

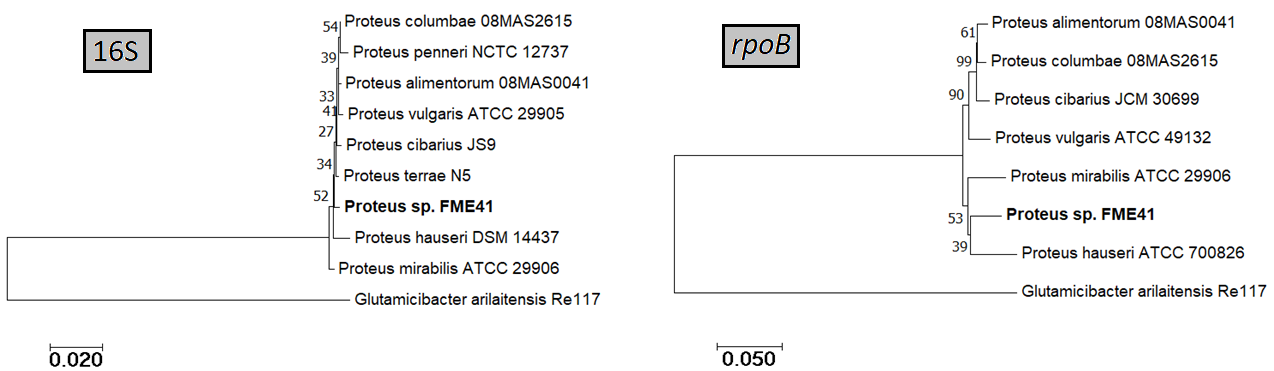

**Figure E.** Phylogenetic tree of *Proteus* genus based on 16S rRNA and *rpoB* gene alignments, including FME41 strain (isolated in this study) and its closest relatives.

In the same way, *Pseudomonas* FME51 strain could not be assigned reliably, althought its 16S rRNA gene sequence shares around 99% identity with the one of *Pseudomonas litoralis* 2SM5^T^, since its *rpoB* nucleotide sequence shares only 93.27% with this type strain (**Table S1, Figure F**).

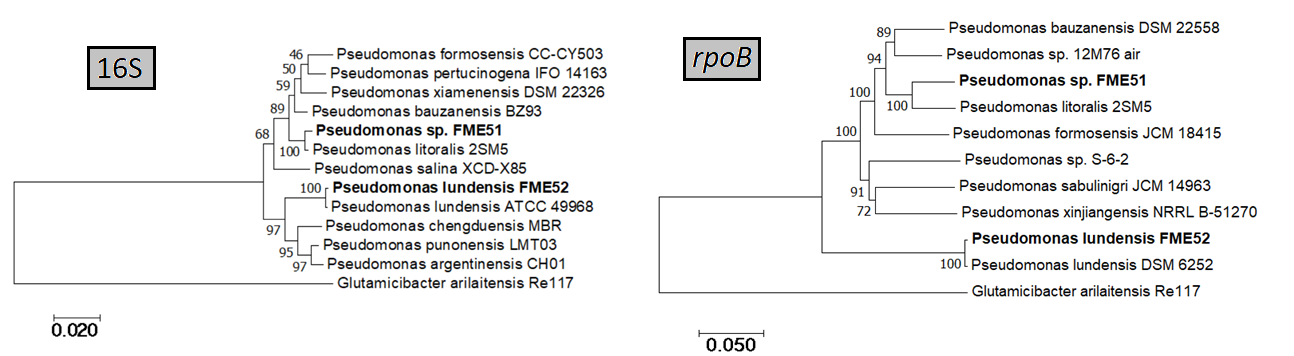

**Figure F.** Phylogenetic tree of *Pseudomonas* genus based on 16S rRNA and *rpoB* gene alignments, including FME51 and FME52 strains (isolated in this study) and their closest relatives.

Furthermore, *Pseudoalteromonas* FME14 and FME53 strains could not be reliably attributed to one species because the two marker genes share high identity (over the threshold) with different species and consequently their result are not congruent (**Table S1, Figure G**).

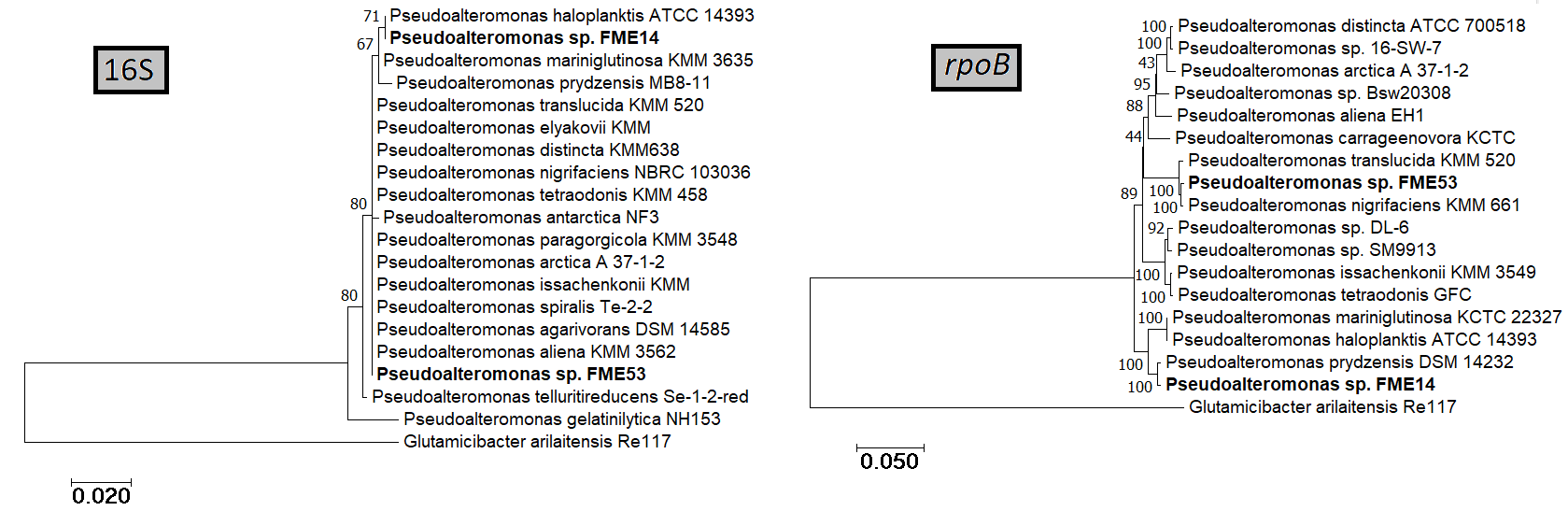

**Figure G.** Phylogenetic tree of *Pseudoalteromonas* genus based on 16S rRNA and *rpoB* gene alignments, including FME14 and FME53 strains (isolated in this study) and their closest relatives.

Finnaly, the four *Psychrobacter* strains remain unassigned because their marker gene sequences display insufficient levels of homology and/or non congruent results to delineate them as defined species (**Table S1, Figure H**).

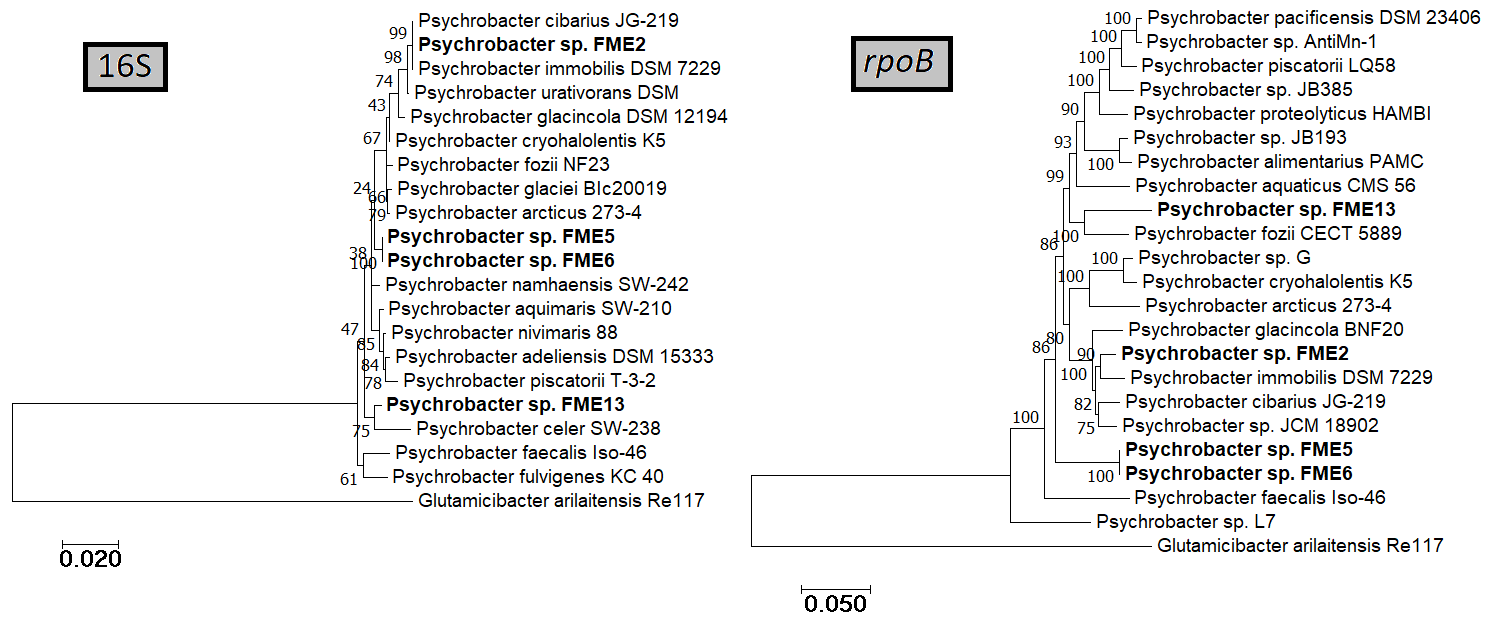

**Figure H.** Phylogenetic tree of *Psychrobacter* genus based on 16S rRNA and *rpoB* gene alignments, including FME2, FME5, FME6 and FME13 strains (isolated in this study) and their closest relatives.
