## Additional File 2 for "Unraveling the world of halophilic and halotolerant bacteria in cheese by combining cultural, genomic and metagenomic approaches"

|  | **Phylum** | **Genus** | **Food sources** | **Main species** | **Potential role or contribution to food** | **Authors** |
| --- | --- | --- | --- | --- | --- | --- |
| **Gram-positive** | Actinobacteria | *Glutamicibacter* | Cheese, milk | *G. arilaitensis, G. bergerei* | Contribution to the colour, flavour and texture properties in cheese or contaminant/ spoilage. | [1-3] |
|  | Actinobacteria | *Brevibacterium* | Cheese, milk, salted food | *B. aurantiacum, B. antiquum, B. linens, B. casei, B.* spp. | Used as adjunct in cheese; proteolytic, lipolytic and esterase activities; production of antmicrobial substances, pigments and aromatic compounds. | [1, 4-10] |
|  | Actinobacteria | *Corynebacterium* | Cheese, milk | *C. casei, C. variabile* | Contribution to the aroma and colour of surface-ripened cheeses. | [11-15] |
|  | Actinobacteria | *Brachybacterium* | Fermented food, especially in cheese | *B. tyrofermentans, B. alimentarium* | Contribution to the flavour and colour in cheeses. | [16-18] |
|  | Firmicutes | *Staphylococcus* (coagulase-negative) | Fermented foods, sausages and meat products | *S. equorum*, *S. succinus, S. xylosus* | Used as adjunct in cheese to improve its texture and bring particular flavors, rind coloration, proteolytic and lipolytic activities. | [5, 13, 19-22] |
|  | Firmicutes | *Oceanobacillus* | Korean fermented food | *O. kimchii, O. gochujangensis* | Unknown function in food, likely derived from solar salts. | [23-25] |
|  | Firmicutes | *Carnobacterium* | Cheese, milk, meat, fish, shrimp | *C. maltaromaticum, C. mobile, C. divergens* | Production of antimicrobial peptides and bacteriocins and production of volatile compounds that contribute for aroma or spoilage in food. | [26-29] |
|  | Firmicutes | *Marinilactibacillus* | Cheese rinds, salted food, cheese brine | *M. psychrotolerans* | Unknown function in cheese, associated to spoilage in dry-cured hams. | [30-33] |
| **Gram-negative** | Proteobacteria | *Advenella* | Cheese, milk | *A. incenata, A. kashmirensis, A.* spp. | Potential to contribute positively to cheese ripening. | [34-36] |
|  | Proteobacteria | *Pseudoalteromonas* | Cheese, Korean fermented food | *P. haloplanktis, P.* spp | Suggestion of play key roles in cheese rind microbial communities. | [37-40] |
|  | Proteobacteria | *Hafnia* | Cheese, milk | *H. alvei* | Used as adjunct culture in cheese to improve cheese flavor. | [29, 39, 41, 42] |
|  | Proteobacteria | *Proteus* | Cheese, milk | *P. vulgaris, P. mirabilis, P. hauseri, P.* spp | Technological interest as contribution to the organoleptic properties of cheese, but also spoiler or potential pathogen. | [43-47] |
|  | Proteobacteria | *Halomonas* | Cheese, milk, meat products | *H. alkaliphila, H. variabilis, H. venusta, H.* spp. | Suggestions of positive impact during cheese ripening or spoilage bacteria. | [14, 34, 37, 48-50] |
|  | Proteobacteria | *Psychrobacter* | Cheese, sea food, meat products | *P. celer, P. cibarius, P.* spp. | Production of volatile compounds could have a positive impact during the cheese ripening or negative impact as in the food spoilage. | [5, 47, 50-52] |
|  | Proteobacteria | *Pseudomonas* | Meat, poultry, milk and dairy products | *P. aeruginosa, P. lundensis, P. fluorescens, P.* spp. | Strongly proteolytic activity that cause undesirable odor and flavors, frequently associated as contaminant or spoilage. | [53-57] |
|  | Proteobacteria | *Vibrio* | Cheese, seafood | *V. casei, V. litoralis, V.* spp. | Suggestion of play a role in the cheese ripening process. | [14, 30, 58-60] |

**Additional File 2. Main bacterial species of halophilic and halotolerant in food and their potential roles.**

1.	Cogan, T.M., *Bacteria, Beneficial: Brevibacterium linens, Brevibacterium aurantiacum and Other Smear Microorganisms☆*, in *Reference Module in Food Science*. 2016, Elsevier.
